# Therapeutic Potential of *Ocimum basilicum* in Diabetes–Malaria Co-morbidity: Evidence from Parasitemia, Biochemical, and Coagulation Outcomes in Mice

**DOI:** 10.1101/2025.09.13.676048

**Authors:** O.O. Ayodele, O.R. Oyerinde, O.N. Bello, D.G. Adeigbe, G.A. Adegbite, F.J. Femi-Olabisi

## Abstract

Diabetes and malaria are major morbidities that impair hepatic and renal functions. This study investigated the effects of the hydromethanol extract of *Ocimum basilicum* (OCB) on parasitemia, biochemical markers, and coagulation parameters in diabetic BALB/C mice infected with *Plasmodium berghei*.

Fifty-six male mice were divided into eight groups (n=7): normal control, diabetes only, malaria only, diabetes + malaria, malaria + OCB, diabetes + OCB, diabetes + malaria + OCB, and diabetes + malaria + metformin. Diabetes was induced by streptozotocin (40 mg/kg, i.p.) for 5 consecutive days, while *Plasmodium berghei* was inoculated in the malaria groups, and infection was confirmed by Giemsa-stained thin smears. Treatments consisted of OCB (100 mg/kg) or metformin (250 mg/kg) orally for 7 days; controls received phosphate-buffered saline.

OCB significantly (p<0.05) reduced parasitemia in infected groups compared with untreated controls. In diabetic and malaria-induced mice, elevated fasting blood glucose, creatinine, and urea were markedly reduced by OCB, with decreases of 33.22%, 70.58%, and 26.32%, respectively, in the malaria + diabetes + OCB group relative to the untreated group. Serum alanine and aspartate aminotransferases were also lowered by OCB more effectively than metformin, indicating hepatoprotective activity. Coagulation profiles showed no significant differences in activated partial thromboplastin time and prothrombin time between OCB-treated and control groups, although prothrombin time decreased in the diabetes + OCB group.

These findings demonstrate that *O. basilicum* possesses anti-plasmodial, antihyperglycemic, and organ-protective effects, highlighting its potential as a source of phytopharmaceutical agents for the treatment and management of malaria and diabetes.

## Introduction

Diabetes mellitus (DM) and malaria are two major global health concerns that often coexist in regions with high burdens of infectious and metabolic diseases. DM, a chronic disorder characterized by persistent hyperglycemia due to impaired insulin secretion or action, is associated with progressive organ dysfunction, including hepatic and renal injury (American Diabetes Association, 2022). Malaria, caused by *Plasmodium* species, remains a leading cause of morbidity and mortality in tropical and subtropical regions, particularly in sub-Saharan Africa (World Health Organization [WHO], 2023). The overlap of these conditions poses a significant public health challenge, as malaria infection can worsen glycemic control, while diabetes may increase susceptibility to infection and complications (Awungafac et al. 2025; Udoh et al., 2020).

The coexistence of diabetes and malaria creates a complex pathophysiological interplay. Hyperglycemia and oxidative stress in diabetes may exacerbate malaria severity, whereas malaria-induced inflammation, hemolysis, and hepatic injury can further impair metabolic regulation (Awungafac et al. 2025; Faure, 2014; Raghunath, 2017). Although conventional therapies such as metformin for diabetes and artemisinin-based combination therapies for malaria are effective, they may be limited by adverse effects, cost, and emerging drug resistance (Bailey, 2024; Li et al., 2025; Rosenthal, 2021). This highlights the need for alternative therapies that are safe, affordable, and capable of addressing multiple pathological targets.

Medicinal plants are widely explored for their phytotherapeutic potential against chronic and infectious diseases. *Ocimum basilicum* (sweet basil), an aromatic herb commonly used in African and Asian traditional medicine, is reported to possess antimicrobial, anti-inflammatory, antioxidant, and hypoglycemic properties (Hussain et al., 2008; Zhakipbekov et al., 2024). Phytochemical studies reveal that it contains phenolics, flavonoids, and essential oils such as eugenol and linalool, which may underlie its pharmacological activities (Dam et al., 2023; Hamid et al., 2024). Despite these promising bioactivities, little is known about its efficacy in the context of diabetes–malaria co-morbidity, particularly regarding parasitemia, biochemical alterations, and coagulation parameters.

This study, therefore, investigated the therapeutic potential of the hydromethanol extract of *O. basilicum* in BALB/C mice with streptozotocin-induced diabetes and *Plasmodium berghei* infection. Specifically, the effects of the extract were evaluated on parasitemia, blood glucose, biochemical indices of liver and kidney function, and coagulation profiles, providing evidence for its possible use as a phytopharmaceutical agent in managing diabetes–malaria co-morbidity.

## Methodology

### Materials and Reagents

Streptozotocin (Santa Cruz, USA), metformin (Glucophage, Merck), methanol, phosphate-buffered, tri-sodium citrate, sodium hydroxide, picric acid, and Giemsa stain were obtained from standard suppliers. Diagnostic assay kits for creatinine, urea, aspartate aminotransferase (AST), and alanine aminotransferase (ALT) were purchased from Randox Laboratories Ltd. Fasting blood glucose (FBG) was determined using Accu-Chek and Fine Test glucometers with compatible strips. Other standard laboratory equipment included a UV–Visible spectrophotometer, centrifuge, rotary evaporator, analytical balance, micropipettes, and glassware.

### Study Design and Animal Handling

Fifty-six (56) male BALB/C mice were procured from the Mountain Top University animal house (Ogun State, Nigeria). Animals were acclimatized for two weeks under standard housing conditions (12 h light/dark cycle, room temperature, free access to food and water). All experimental procedures followed the Institutional Animal Ethics Committee (IAEC) guidelines. Institutional ethic approval was obtained with reference number CMUL/ACUREC/09/22/926.

The mice were randomly assigned into eight groups (n = 7):

1. Normal control
2. Diabetes only
3. Malaria only
4. Diabetes + malaria
5. Malaria + *O. basilicum* extract
6. Diabetes + *O. basilicum* extract
7. Diabetes + malaria + *O. basilicum* extract
8. Diabetes + malaria + metformin

### Collection and Extraction of Plant Material

Fresh *Ocimum basilicum* was collected in April 2023 from Ibafo, Ogun State, Nigeria, and authenticated by Dr. Nodza George (Department of Botany, University of Lagos). A voucher specimen (No. 10040) was deposited at the University herbarium. Leaves and roots were air-dried at 40 °C, pulverized, and stored in airtight containers at 4 °C.

For extraction, 50 g of powdered leaves were macerated in 200 mL methanol and 200 mL distilled water (8:1 v/w) for 120 h with intermittent shaking. The filtrate was concentrated using a rotary evaporator and dried in a hot-air oven at 50 °C to yield a semi-solid extract, which was stored at 4 °C until use.

### Induction of Diabetes and Malaria

Diabetes was induced by intraperitoneal administration of STZ (40 mg/kg) for five consecutive days following an overnight fast (Furman, 2021). Hyperglycemia was confirmed on day 14 in mice with FBG > 190 mg/dL.

Malaria infection was induced with *Plasmodium berghei* (NK65 strain) obtained from the Institute of Advanced Medical Research and Training, University of Ibadan. On day 9, diabetic groups were inoculated intraperitoneally with 0.2 mL of parasitized blood diluted in saline. Parasitemia was monitored by Giemsa-stained thin blood smears from tail snips.

### Treatment Protocol

OCB extract (100 mg/kg) and metformin (250 mg/kg) were administered orally for seven consecutive days. Normal control groups received PBS.

### Sample Collection

On day 25, animals were anesthetized and sacrificed by cervical dislocation. Blood was collected via ocular puncture into tubes with 3.2% trisodium citrate for coagulation assays. Plasma was separated by centrifugation (2500 g, 10 min). Liver and kidney tissues were excised, homogenized in PBS (1:8 w/v), centrifuged (12,000 g, 10 min), and supernatants stored at –20 °C.

### Biochemical Assays

ALT and AST activities were determined using Randox kits based on 2,4-dinitrophenylhydrazine methods. Serum creatinine and urea were quantified enzymatically using Randox kits, while total protein was determined by the Biuret method.

### Hematological and Coagulation Parameters

Packed cell volume (PCV) was measured using the microhematocrit method. Prothrombin time (PT) and activated partial thromboplastin time (aPTT) were determined using standard coagulation kits in accordance with the manufacturer’s instructions.

### Statistical Analysis

Data were expressed as mean ± SEM. One-way ANOVA followed by Tukey’s post-hoc test was performed using GraphPad Prism (v9.0.0.121). Significance was set at p < 0.05.

## Results

### Effects of *Ocimum basilicum* on Fasting Blood Glucose (FBG)

Diabetic mice exhibited a significant elevation in FBG levels on Days 15 and 25 compared to the normal control. Treatment with OCB extract significantly reduced FBG in diabetic (Db+OCB), malaria-diabetic (Mal+Db+OCB), and malaria-only (Mal+OCB) groups relative to their untreated counterparts (Figure 1).

**Figure 1.**
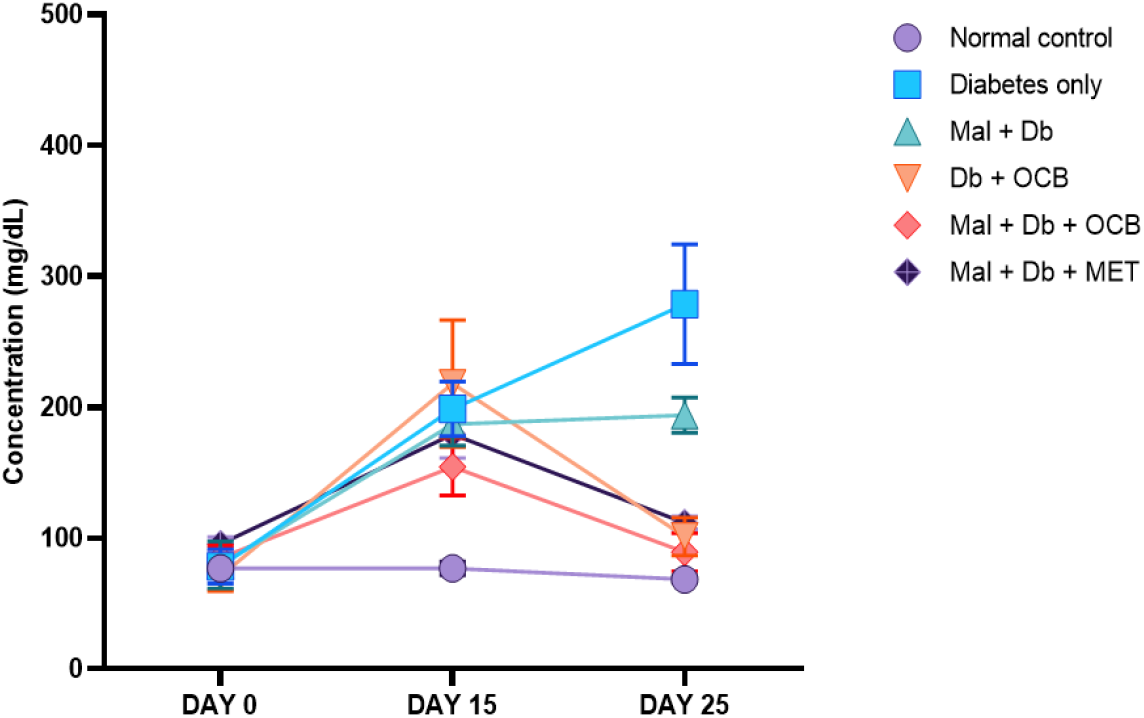
Effect of OCB on FBG levels in experimental mice. MAL -Malaria, Db - Diabetic, OCB - *Ocimum basilicum*, MET - Metformin.

### Effects of *O. basilicum* on Parasitemia

Parasitemia was consistently highest in the malaria-only group throughout the study. OCB-treated mice (Mal+OCB, Mal+Db+OCB) showed a reduced parasitemia burden, particularly from Day 9 onwards, compared to untreated controls. Metformin also decreased parasitemia in Mal+Db mice at Day 25 (Figure 2).

**Figure 2.**
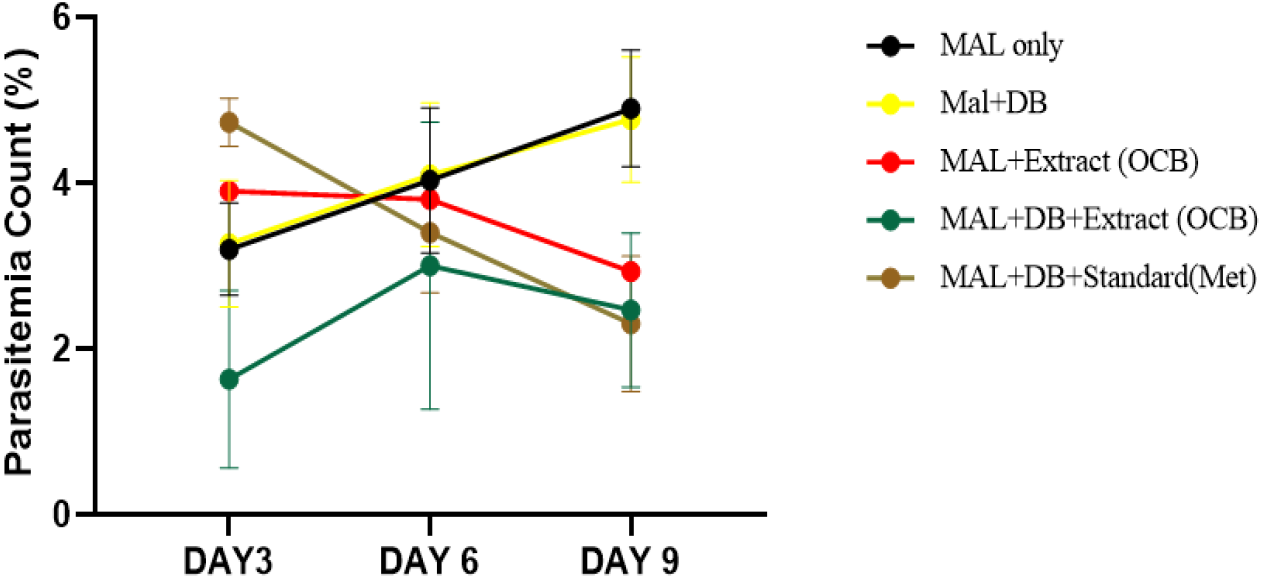
Effect of OCB on parasitemia count in experimental mice. NC-Normal control, MAL -Malaria, DB - Diabetic, OCB - *Ocimum basilicum*, Met - Metformin.

### Effects of *O. basilicum* on Body Weight

Infection with *P. berghei* and/or diabetes induced progressive weight loss. However, OCB treatment attenuated this decline, with treated groups maintaining higher body weights compared to their respective controls. By Day 25, weight preservation was most evident in Db+OCB and Mal+Db+OCB groups (Figure 3).

**Figure 3.**
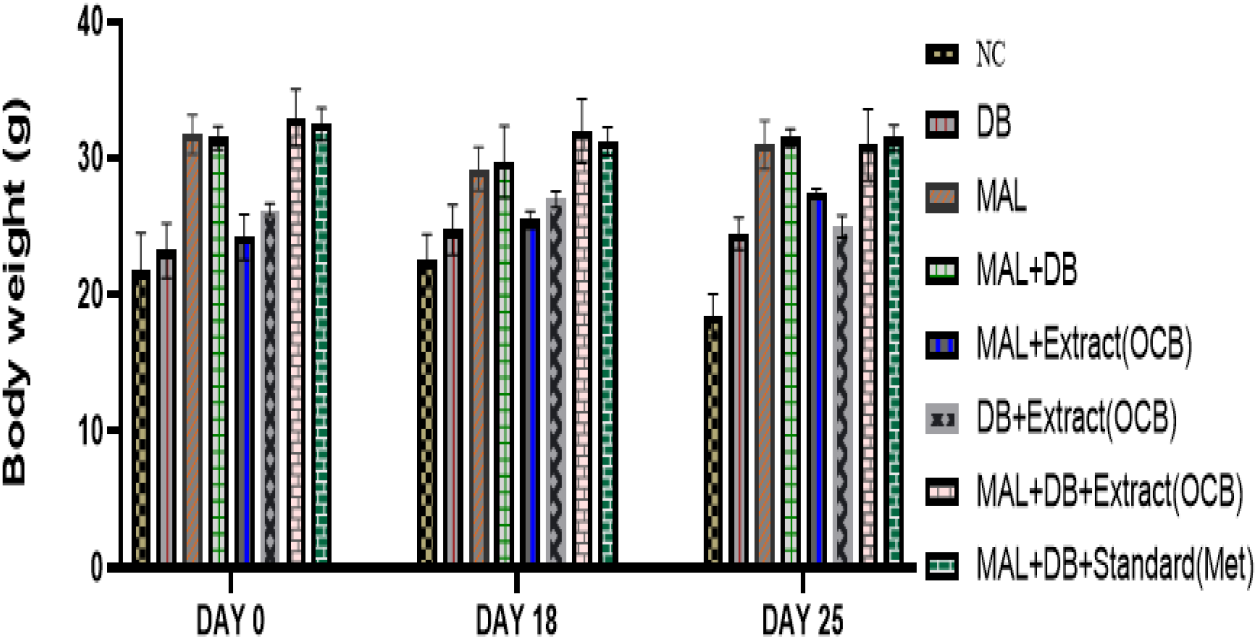
Effect of OCB on the body weight of experimental mice. NC-Normal control, MAL -Malaria, DB - Diabetic, OCB - *Ocimum basilicum*, Met - Metformin.

### Effects on Liver Enzyme Activities (ALT and AST)

ALT levels were significantly elevated in untreated disease groups (Db, Mal, Mal+Db) compared to the normal control. Treatment with OCB or metformin attenuated these elevations (Figure 4). AST concentrations were significantly reduced in Db+OCB compared to Db-only mice. Conversely, Mal+Db mice showed higher AST activity than Mal-only or Db-only groups, while OCB treatment significantly reduced AST compared to untreated malaria groups (Figure 5).

**Figure 4.**
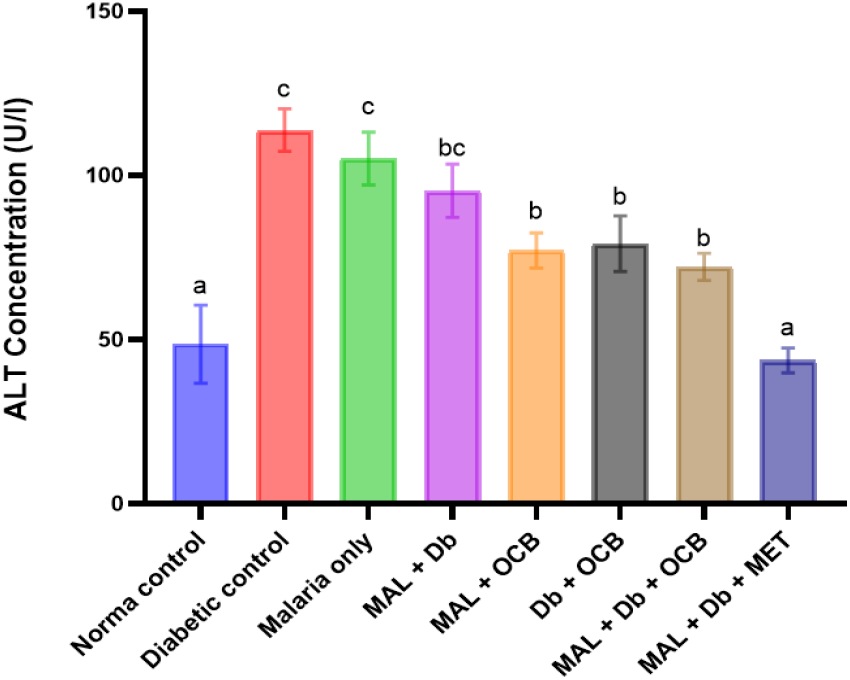
Effect of OCB on ALT concentrations in experimental mice. Data are presented as Mean ± SEM; n=6. Different letters on bars indicate a significant difference (p<0.05). MAL -Malaria, Db - Diabetic, OCB - *Ocimum basilicum*, MET - Metformin.

**Figure 5.**
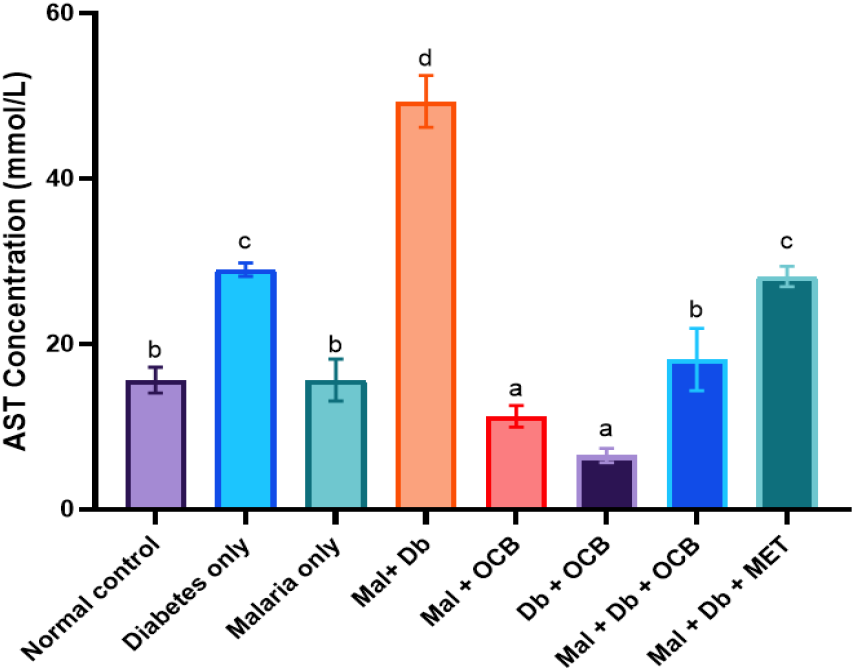
Effect of OCB on AST concentrations in experimental mice. Data are presented as Mean ± SEM; n=6. Different letters on bars indicate a significant difference (p<0.05). Mal -Malaria, Db - Diabetic, OCB - *Ocimum basilicum*, MET - Metformin.

### Effects on Renal Function Markers (Urea and Creatinine)

Urea levels were elevated in Mal-only mice but significantly reduced in Mal+OCB (Figure 6). Creatinine concentrations were significantly reduced in OCB-treated groups (Mal+OCB, Db+OCB, Mal+Db+OCB) compared to untreated controls. The creatinine concentrations were significantly decreased in treatment groups (Mal + OCB, Db + OCB, and Mal +Db +OCB) when compared to normal control (Figure 7).

**Figure 6.**
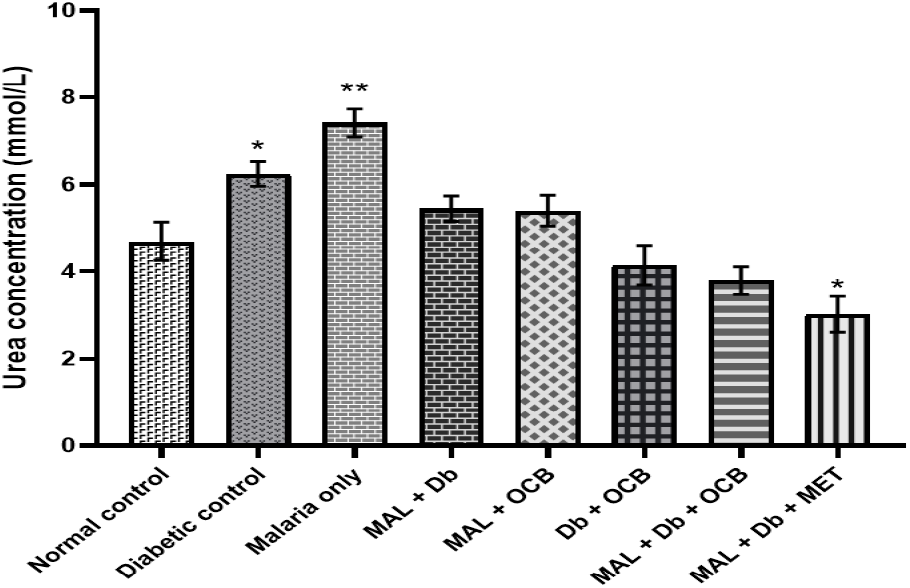
Effect of OCB on Urea concentrations in the mice. Data are presented as Mean ± SEM; n=6; *p<0.05, **p<0.01 compared to corresponding OCB-treated groups. Mal -Malaria, Db - Diabetic, OCB - *Ocimum basilicum*, MET - Metformin.

**Figure 7.**
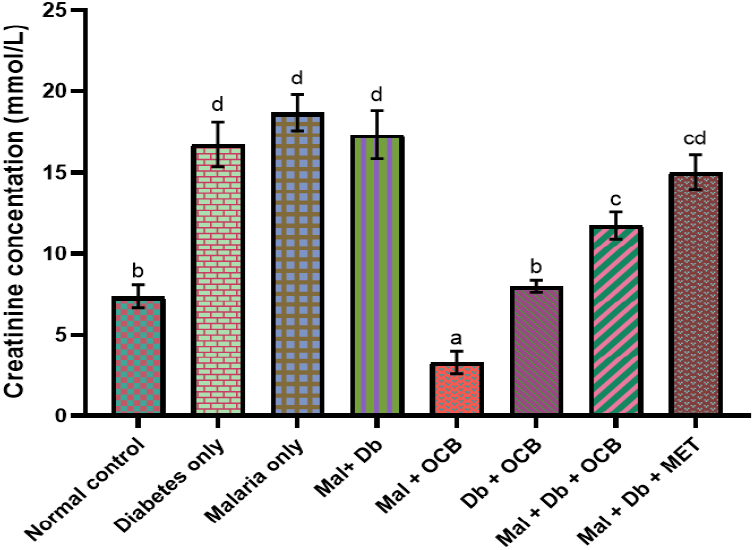
Effect of OCB on Creatinine concentrations in the mice. Data are presented as Mean ± SEM; n=6. Different letters on bars indicate a significant difference (p<0.05). Mal -Malaria, Db - Diabetic, OCB - *Ocimum basilicum*, MET - Metformin.

### Effects on Packed Cell Volume (PCV)

At Day 25, PCV values were significantly higher in Mal+Db+OCB and Mal+Db+Met groups compared with untreated Mal+Db mice, suggesting an anti-anemic effect of OCB (Figure 8).

**Figure 8.**
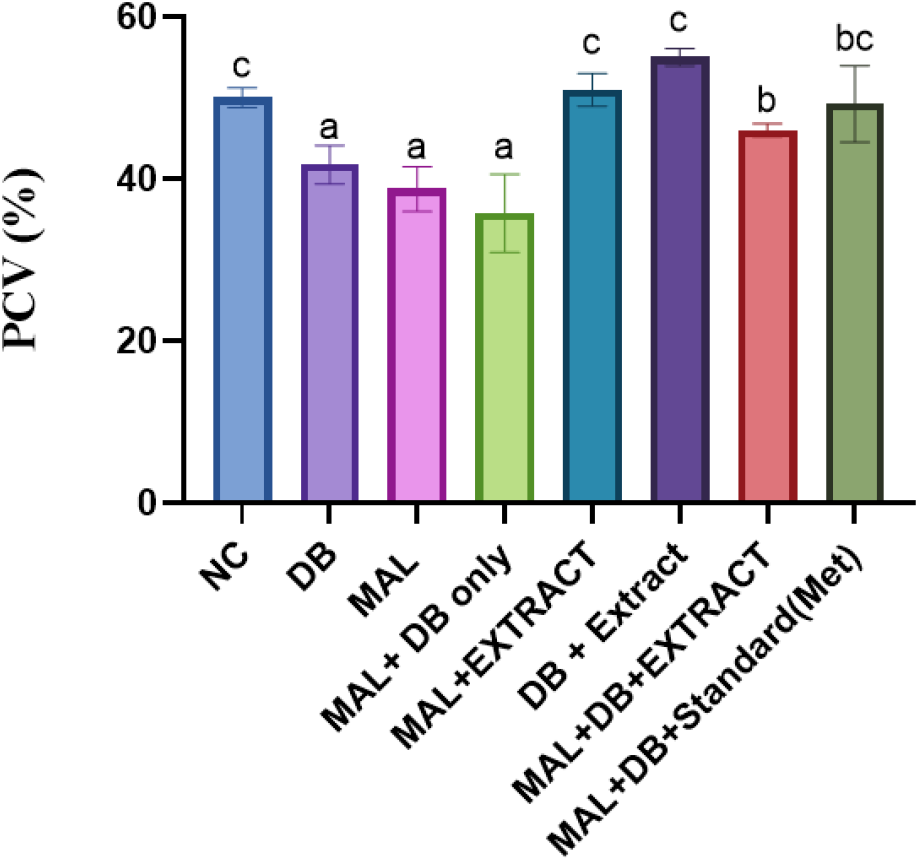
Packed Cell Volume (%) of experimental mice on day 25. NC- Normal control, MAL -Malaria, DB - Diabetic, OCB - *Ocimum basilicum*, Met - Metformin.

### Effects on Coagulation Parameters (Activated Partial Thromboplastin Time; APTT) and Prothrombin Time; PT)

The APTT of normal controls was significantly longer than all experimental groups. OCB shortened APTT in malaria-infected mice compared to Mal-only, suggesting a pro-coagulant effect (Figure 9). PT was shorter in the Db-only group compared to controls, while OCB did not significantly alter PT across most groups, except Db+OCB (Figure 10).

**Figure 9.**
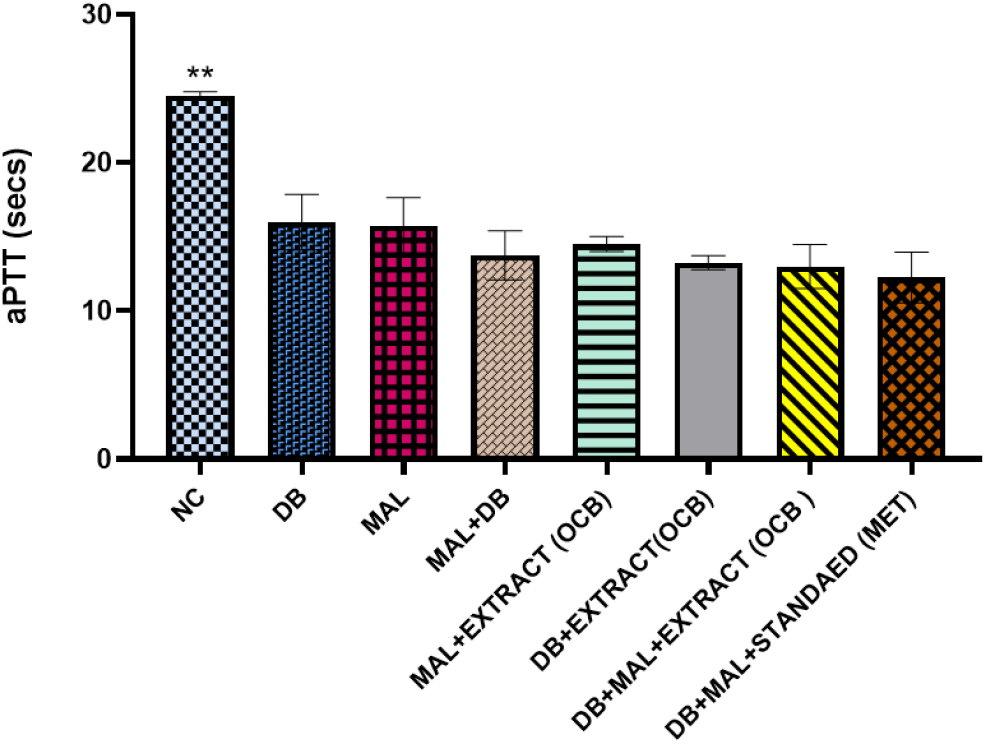
Activated Partial Thromboplastin Time of experimental mice. Values represented as Mean ±SEM, n=6, **p<0.01 significantly higher than other groups. NC-Normal control, MAL -Malaria, DB - Diabetic, OCB - *Ocimum basilicum*, MET - Metformin.

**Figure 10.**
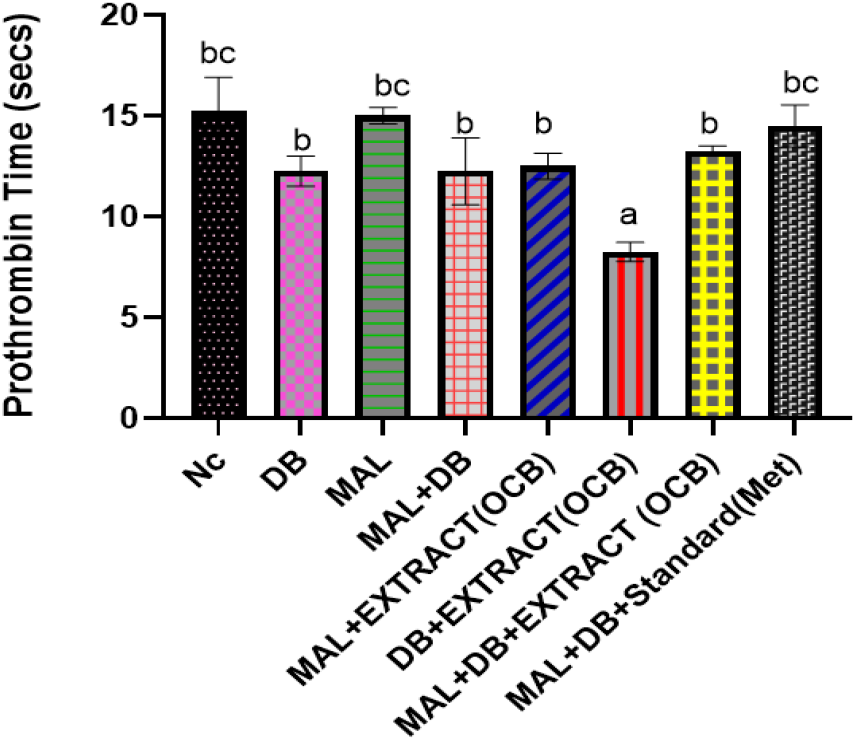
Prothrombin Time of experimental mice. Values represented as Mean ±SEM, n=6, Different letters on bars indicate a significant difference (p<0.05). NC-Normal control, MAL -Malaria, DB - Diabetic, OCB - *Ocimum basilicum*, MET - Metformin.

### Plasma and Liver Total Protein (TP)

Plasma TP was significantly reduced in Mal+Db compared to normal controls. Treatment with OCB modestly improved TP levels, while liver TP was significantly elevated in malaria-treated groups compared to controls (Figure 11).

**Figure 11.**
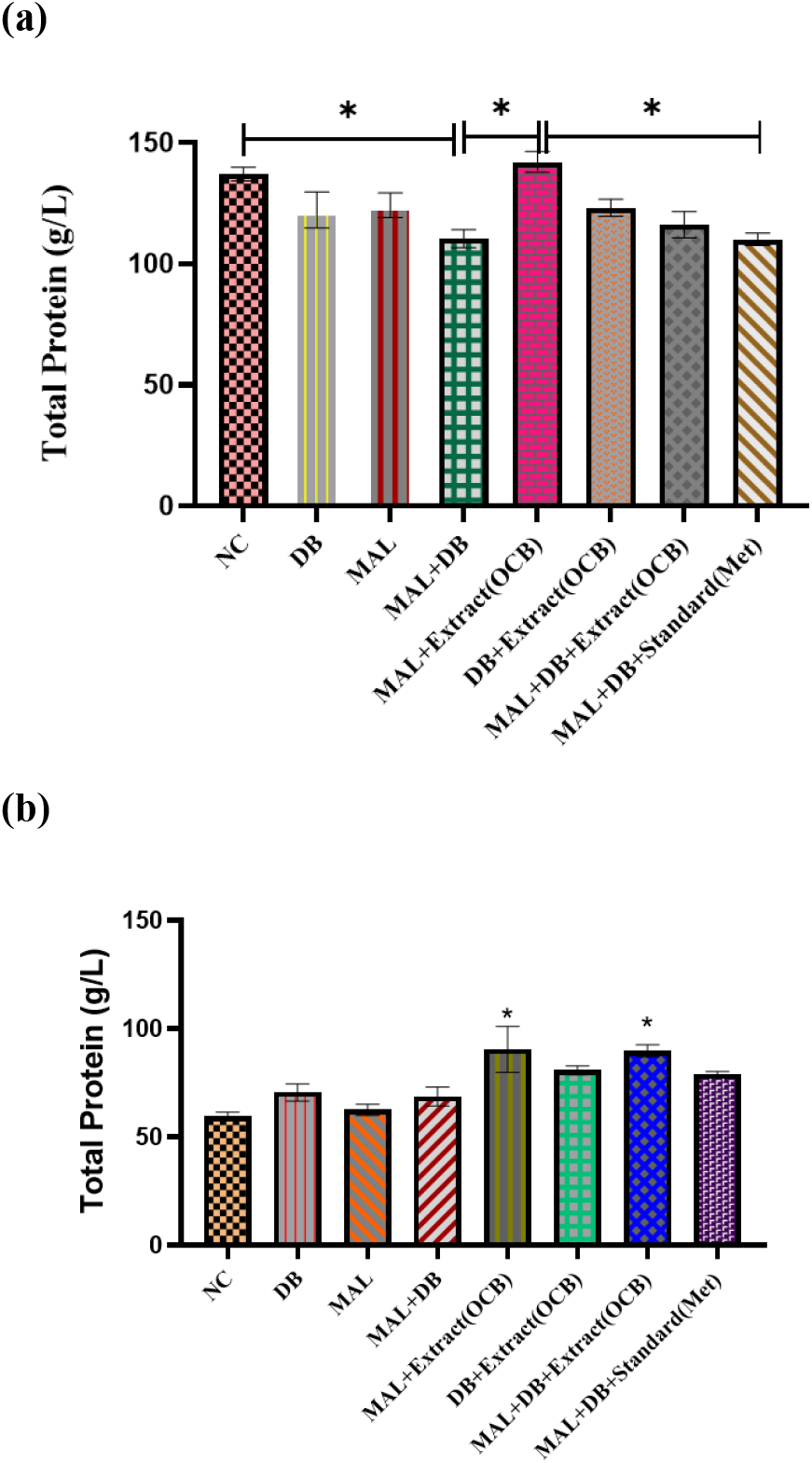
Plasma (a) and Liver (b)Total Protein concentrations of experimental mice. Values represented as Mean ±SEM, n=6, *p<0.0 significantly different. NC-Normal control, MAL -Malaria, DB - Diabetic, OCB - *Ocimum basilicum*, MET - Metformin.

## Discussion

The present study investigated the therapeutic potential of *Ocimum basilicum* (OCB) methanol extract in diabetes–malaria co-morbidity, focusing on parasitemia, biochemical, and coagulation outcomes in mice. The findings demonstrate that OCB exhibits hypoglycemic, antiplasmodial, organ-protective, and hematological benefits, supporting its potential role in managing comorbid metabolic and infectious diseases.

The induction of diabetes significantly elevated FBG levels in diabetic mice on Days 15 and 25 compared with the normal control, confirming the hyperglycemic state. Treatment with *Ocimum basilicum* (OCB) markedly reduced blood glucose concentrations in diabetic mice, as well as in the comorbid groups (Mal + Db + OCB and Mal + OCB). This hypoglycemic effect is consistent with previous reports that *O. basilicum* possesses antihyperglycemic properties through modulation of carbohydrate metabolism and enhancement of insulin sensitivity (El-Beshbishy et al., 2012; Yedjou et al., 2023). The glucose-lowering ability of OCB may be attributed to its flavonoids, terpenoids, and phenolic compounds, which have been shown to enhance pancreatic β-cell activity, improve glucose uptake in peripheral tissues, and inhibit α-glucosidase activity (Javanmardi et al., 2003; Singh et al., 2022).

Parasitemia was highest in the malaria-only group, confirming a successful infection. OCB treatment significantly lowered parasitemia in both malaria-infected and comorbid groups, especially by Day 9, indicating an anti-plasmodial effect. The decrease was similar, though not identical, to that seen with metformin treatment. The anti-parasitic activity might be due to phytochemicals in *O. basilicum* such as eugenol, linalool, and rosmarinic acid, which have been shown to interfere with Plasmodium growth and replication (Bunalema et al., 2025; Chinsembu, 2015; Chukwuma et al., 2023). Additionally, better glycemic control in diabetic mice could improve immune responses against malaria, thereby lowering parasitemia.

Weight loss is a common feature in both malaria and diabetes due to increased catabolism, anorexia, and impaired nutrient utilization. The OCB-treated groups exhibited attenuated body weight loss compared to untreated controls. In fact, OCB promoted weight gain in both single-disease and comorbid conditions. This finding suggests a protective effect on energy balance, possibly by reducing oxidative stress and improving metabolic efficiency. Flavonoid-rich plant extracts, including OCB, are known to preserve muscle protein and fat stores during metabolic stress (Jomova et al., 2025; Ullah et al., 2020).

The malaria-diabetic mice treated with OCB and those treated with metformin showed significantly higher PCV than the untreated comorbid group. Since malaria typically induces hemolytic anemia, the restoration of PCV indicates that OCB exerts a hematoprotective effect, possibly through antioxidant activity and inhibition of hemolysis. Phytochemicals such as rosmarinic acid and eugenol are known to stabilize erythrocyte membranes and reduce oxidative damage, thereby mitigating anemia (Magtalas et al., 2023; Tsuchiya et al., 2015; Ydyrys et al., 2023).

ALT and AST are markers of liver cell integrity. In this study, ALT levels were significantly increased in the negative controls (Diabetes only, Malaria only, and Mal + Db), indicating liver injury caused by hyperglycemia, oxidative stress, and parasitemia. Interestingly, ALT levels were also elevated in OCB-treated groups, suggesting either a temporary liver stress response or increased liver enzyme activity. However, these levels remained below those of the negative controls. Conversely, AST levels decreased in Db + OCB compared to diabetic-only mice, while malaria-diabetic mice showed the highest AST levels. Treatment with OCB reduced AST levels in malaria groups. These results indicate that OCB offers some hepatoprotective effects, especially against malaria-related damage (Meng et al., 2020; Osano et al., 2024).

Urea levels were significantly higher in the malaria-only group, consistent with renal dysfunction due to malaria-associated hemolysis and dehydration. Treatment with OCB significantly reduced urea levels, suggesting nephroprotection. Similarly, creatinine levels were decreased in OCB-treated diabetic and malaria groups compared with negative controls, and also lower than the normal control. This pattern indicates that OCB may protect against renal impairment induced by diabetes and malaria. Flavonoids and polyphenols in OCB likely reduce oxidative and inflammatory stress in renal tissues, thus improving kidney function (Anwar et al., 2021; Félix et al., 2018).

In malaria mice, OCB shortened APTT relative to untreated controls, indicating a possible procoagulant effect. This effect could be beneficial in malaria, where prolonged clotting times due to thrombocytopenia and coagulation factor depletion are common (Akinosun et al., 2017; Ghosh and Shetty, 2008; Tesfaye et al., 2024). In diabetic mice, PT was shortened relative to normal controls, reflecting a hypercoagulable state. OCB treatment did not significantly alter PT across groups, except in the diabetic-treated group, suggesting a mild normalization effect. Taken together, these results imply that OCB may modulate coagulation pathways differently depending on the pathological context, potentially balancing bleeding and clotting risks.

Plasma TP levels were significantly reduced in the Mal + Db group compared to normal control, consistent with impaired protein synthesis and catabolism during chronic infection and hyperglycemia. Treatment with OCB did not markedly alter plasma TP except in the malaria-treated group, which showed improvement. Interestingly, liver TP was significantly increased in malaria-treated groups compared to controls, suggesting enhanced hepatic protein synthesis. This may indicate hepatostimulatory properties of OCB, supporting improved recovery from malaria-induced hepatic dysfunction (Nnamudi et al., 2020).

## Conclusion

The findings show that *Ocimum basilicum* provides multiple health benefits in experimental models of diabetes, malaria, and their co-occurrence. These benefits include antihyperglycemic, antiplasmodial, hematoprotective, nephroprotective, and moderate hepatoprotective effects, as well as effects on coagulation and protein metabolism. Overall, these results suggest that *O. basilicum* is a promising phytopharmacological source for treating and managing malaria and diabetes, though further mechanistic and toxicological studies are needed for clinical translation.

## Conflict of Interest

The authors declare no conflict of interest.

## Authors’ Declaration

The authors hereby declare that the work presented in this article is original.

## Acknowledgments

The authors would like to thank Mr. Opeyemi Ojo of the Department of Chemical Sciences and Mr.Gabriel Abah of the Department of Biochemistry, both at Mountain Top University, Nigeria, for their technical support.

## Notes

### Competing Interest Statement

The authors have declared no competing interest.

